# A recurrent neuronal model for the effect of predictions on sensory processes

**DOI:** 10.1101/2021.02.05.429913

**Authors:** Buse M. Urgen, Hilal I. Basturk, Huseyin Boyaci

## Abstract

The effects of prior knowledge, predictions, and expectations on sensory and decision-making processes have been extensively studied. Yet, the neural mechanisms underlying those effects are still unclear. Here, we propose a recurrent neuronal model and test its predictions on behavioral and neuroimaging data from the literature. The model implements predictive processing through recurrent interactions among feature-tuned neural populations that integrate feedback and feedforward signals within established cortical circuitry. Our results show that the model can successfully explain the behavioral effects of prediction found in a previous study in which houses and faces were used as stimuli. We then simulate fMRI data using the protocols of three different studies from the literature, and the optimized model parameters obtained from the model fit to the behavioral data. Although the studies used diverse visual stimuli, not limited to faces and houses, the model predicts their findings to a great extent, proving its generalizability. Overall, our findings demonstrate that the proposed model can provide a link between behavior and neural activity, offering a theoretical account of how prior knowledge, predictions, and expectations influence sensory processing.

## Introduction

A growing body of work over the past two decades has examined whether and how predictions, prior knowledge, and expectations affect sensory processes (e.g., de Lange et al., 2018). These studies have consistently shown that predicted stimuli are detected more rapidly and accurately than unpredicted stimuli. While these effects are well-established at the behavioral level and often shown to be consistent with the Bayesian rule, underlying neural computations remain relatively unclear.

In this framework, priors represent learned regularities of the environment that shape perceptual expectations. These priors can arise from recent experience within an experimental context or from long-term adaptation to environmental statistics. Recent evidence further suggests that such environmental priors may be embedded directly within sensory measurements through the tuning properties of early visual neurons, providing a biologically efficient mechanism for implementing prior expectations (Harrison et al., 2023). Consistent with this, preparatory neural activity has been observed in sensory regions even before stimulus onset, reflecting top-down predictive signals that bias sensory processing in anticipation of expected input (Summerfield & De Lange, 2014). Together, these findings highlight the importance of identifying the neural computations that implement such expectations, and several “predictive processing” models have been proposed for the related brain function (e.g., Friston, 2005; Heeger, 2017; D. Mumford, 1992; Rao & Ballard, 1999). At a common and fundamental level, all predictive processing models suggest that information processing in the brain can be implemented through dynamic interactions between bottom-up sensory input and top-down predictions. Among them, the “predictive coding” model (Friston, 2005; Rao & Ballard, 1999) has gained strong support in the literature. The model proposes that the brain minimizes the prediction error (the difference between actual and predicted sensory input) through hierarchical processing.

While the predictive coding model and its flavors provide one possible mechanism, they are not the only way to conceptualize the interaction between sensory input and predictions. In this study, we propose a recurrent cortical model based on the framework of Heeger (2017) as an alternative, parsimonious account for understanding how predictions shape perceptual processes. Unlike the predictive coding model, the proposed model does not assume or posit distinct sub-populations of neurons for representations and errors (see Discussion). Instead, it proposes that neural activity emerges from the interplay between the weighted influence of bottom-up sensory input and top-down predictions using the neural infrastructure currently known to exist.

To test the model, we used behavioral and neuroimaging results from the literature. First, we modeled the behavioral findings of Urgen and Boyaci (2021), where the authors found that unmet expectations lead to higher temporal thresholds compared to neutral and expected conditions. In that study, each trial began with an informative or neutral cue followed by a briefly presented intact image, and observers indicated whether the intact image appeared on the left or right of a central fixation mark on the screen. Duration thresholds were computed for expected, neutral, and unexpected conditions. Using a recursive Bayesian model, this earlier study showed that higher thresholds can be explained by longer times needed by the system to complete the sensory process when sensory input and expectations disagree. To anticipate, we found that our proposed cortical model can capture the behavioral results and approximate the Bayesian rule at this level. Next, using the model parameters obtained from the fit to the behavioral data, we have simulated the results of three fMRI studies in the literature. In other words, we generated synthetic data as if the participants in the behavioral experiment participated in those fMRI studies.

Critically, a variety of features (i.e., visual stimulus properties such as oriented gratings, moving dots, or object categories) were used in those fMRI experiments, not limited to houses and faces as in the behavioral experiment. This stringent approach allowed us to test the model and its generalizability in an unbiased manner. We present our findings below and discuss their implications, with a particular emphasis on comparing the proposed model to the predictive coding model.

## Methods

### Model

The model consists of one input, one decision, and three intermediate layers, each containing category-specific feature units (representing populations of neurons; e.g., face, house, and scrambled in the behavioral experiment, as described below). We first define an optimization function that the system tries to minimize:

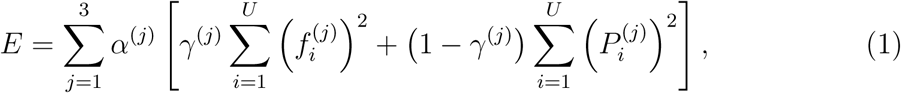

where U is the number of features, indices i and j run over units (features) and layers, respectively, and

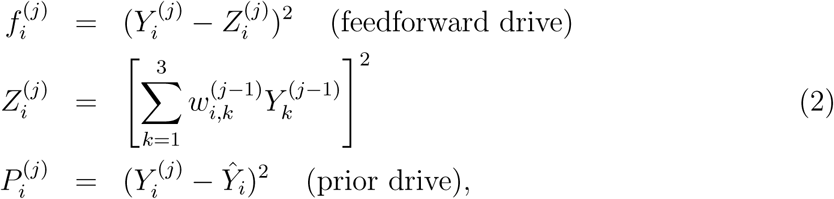

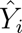 are prior, and 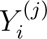 are intermediate layer unit responses. *γ*^(*j*)^ can have values between 0 and 1, and determine the relative weights of the feedforward and prior drives, whereas *α*^(*j*)^ determine the relative contributions of layers to E. 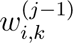 are the weights of connections between units of different layers. Unit responses are updated by minimizing the optimization function with respect to 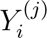 using gradient descent.

Note that feedback and “horizontal” interactions within the same layer emerge after taking the derivative of the optimization function. At the onset of a trial or stimulation (*t* = 0), intermediate layer unit responses are randomly drawn from a normal distribution (Ma et al., 2006).

#### Input layer units

We define the input stimulus, **s**, as a vector with U elements. For example, for the behavioral experiment (see below)

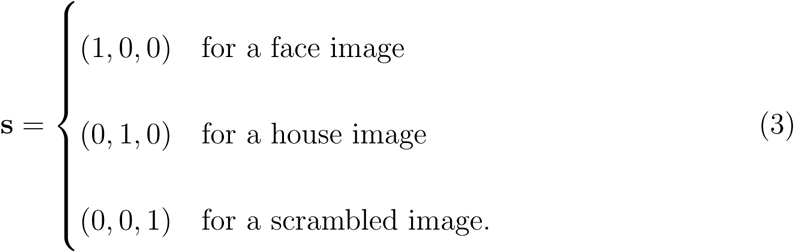

At each iteration, we first compute a noisy, abstracted observation **x** drawn from a normal distribution with mean **s**.

Next, we calculate the input layer responses drawn from a normal distribution based on their hypothetical “tuning” curves

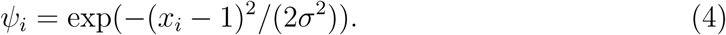

Note that the input layer units are not subject to the optimization, and they do not receive feedback or prior drive.

#### Prior units

We define *initial prior probabilities*, **c** = (*c*_1_,…, *c_U_*), which depend on the cue and its validity at the beginning of a trial or stimulation (t = 0).

Then we initialize the activity of prior units by drawing values from a normal distribution with mean **c**.

For *t* > 0, the prior unit activities are updated recursively at each iteration with the past values of 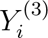 unit responses (the third and last intermediate layer). Specifically, the values of 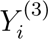 in the previous iteration become the prior in the next iteration.

### Model Fitting to Behavioral Data

We tested whether the cortical model can explain the observed behavioral effect in Urgen and Boyaci (2021), where observers were asked to make a decision about the location (left or right) of a briefly presented intact image, and their duration thresholds were computed (Fig. 1). First, we assumed that computations were carried out iteratively for the duration of a trial. Thus, the number of iterations is

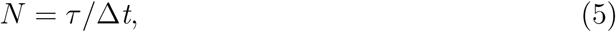

where *τ* is the duration of the presentation of the images in the trial, and Δ*t* determines how long each iteration lasts in the system. Next, the model made a decision based on the sum of the last layer’s (Layer 3) face and house unit responses for left- and right locations (*T_LEFT_*, *T_RIGHT_*)

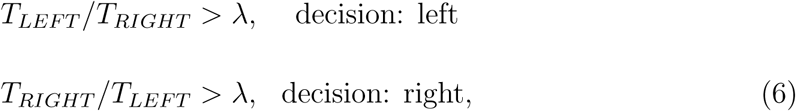

where *λ* is the decision threshold (Heekeren et al., 2004). If the above-mentioned conditions are not satisfied, a choice is made randomly. We fit the model to the observer data at the individual participant level by optimizing three parameters: *λ* (decision criteria), Δ*t* (duration of an iteration), and *σ* (standard deviation of the tuning curves at the input layer). The parameters were optimized with a basic Gaussian mutation genetic algorithm using MATLAB’s Global Optimization Toolbox.

**Figure 1:**
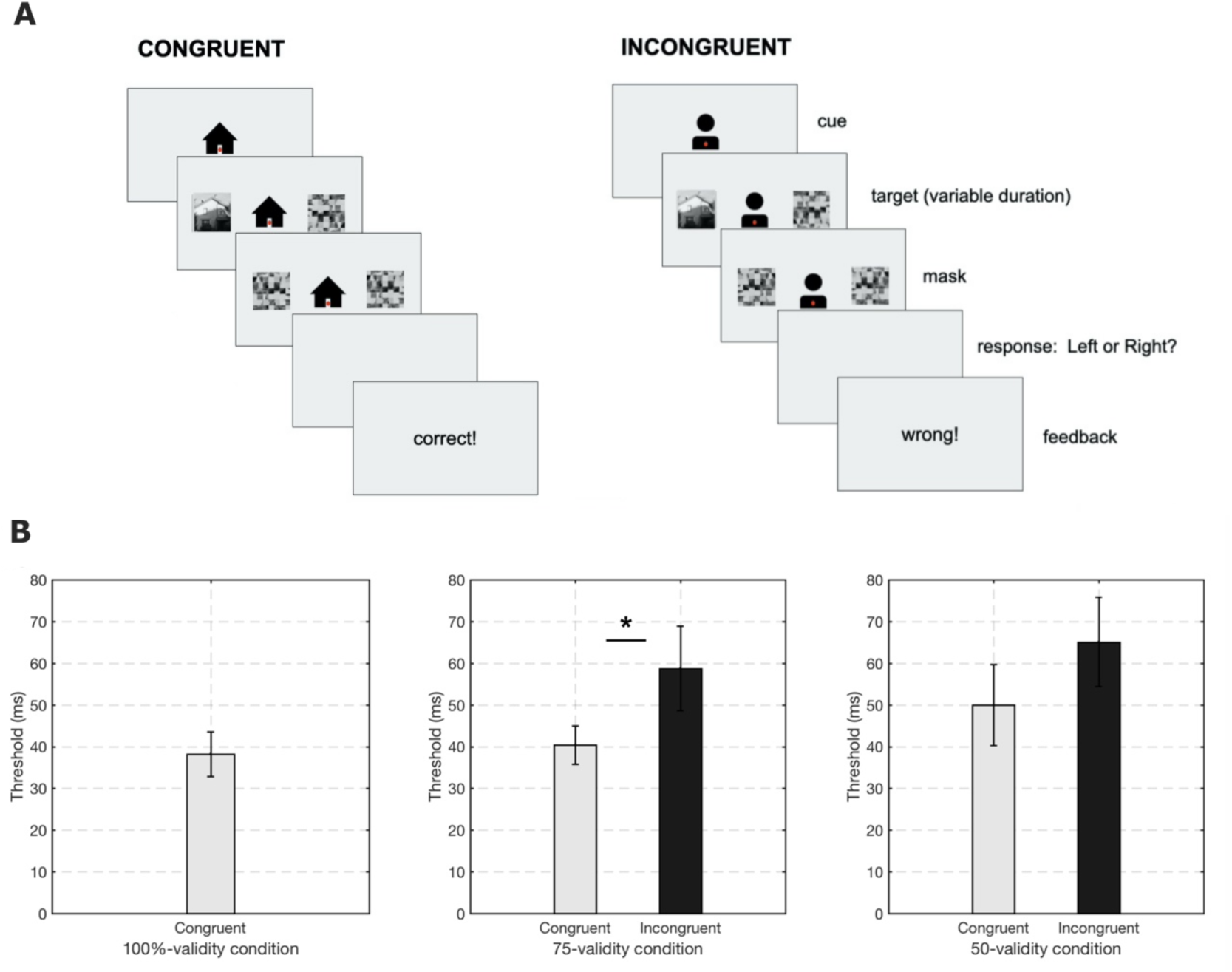
Behavioral experiment (Urgen & Boyaci, 2021). **A. Experimental design**. At each trial, task-irrelevant predictive cues, a face or a house symbol, provided prior information about the probability of the upcoming target image category. Next, an intact target image and a scrambled version of it were presented on the left and right periphery. Presentation duration of the target image varied at each trial and was determined by an adaptive staircase procedure. Participants’ task was to detect the spatial location of the target image, left or right. The validity of the cue was set at 100%, 75%, and 50% in separate sessions. **B. Results**. Bar plots show the temporal thresholds to successfully detect the spatial location of the target image. Incongruent (unpredicted) trials led to higher temporal thresholds than congruent (predicted) trials in the 75%-validity condition. There were no other significant differences. This figure is adapted from the results and figures presented in Urgen and Boyaci (2021). Copyright 2021, Elsevier.

To establish a direct correspondence with the behavioral measures used in the original empirical study, we computed simulated temporal thresholds as follows. First, the number of iterations, *N*, required to reach the temporal threshold accuracy cutoff (77% detection accuracy) was interpolated from the model fits. Then, to convert the measure to the time domain, these were multiplied by each subject’s optimal iteration duration, Δ*t*.

### Neural Data Simulations

To check the biological plausibility and generalizability of the model, we compared model predictions with the results of three fMRI studies in the literature (Egner et al., 2010; Kok et al., 2011; Rahnev et al., 2011).

To this end, we used the optimized parameter values (*λ*, Δ*t*, *σ*) of each participant from Urgen and Boyaci (2021) as explained above, and simulated neural activity following the experimental protocols of the studies.

Next, the simulated neural activity was convolved with a double-gamma hemodynamic response function (HRF), and BOLD responses for specific trials were computed using the beta weights of General Linear Model (GLM) regressors. Finally, these BOLD responses were averaged across participants to obtain a group mean. To compare the simulations with the empirical results, we performed the statistical tests used in the original studies, which included repeated measures ANOVAs and post hoc Student’s *t*-tests.

## Results

### Model Fitting to Behavioral Data

The cortical model we propose here is tested on the behavioral findings of Urgen and Boyaci (2021). Figure 1 shows the experimental paradigm and behavioral results of the study. Briefly, each trial started with a foveally presented cue, which was either a house or a face symbol. (The study also contained a condition where the cue was uninformative, a question mark. Because of the different nature of those trials, they are not included in the model simulations here.) This predictive cue was informative about the upcoming target image category with varying validity set at 100%, 75%, and 50% in different experimental sessions. Subsequently, an intact (target) image (face or house) and its scrambled version were briefly shown on either side of the central fixation point, followed by new scrambled images as masks. Participants’ task was to indicate the location, left or right, of the intact image irrespective of its category. Temporal thresholds were computed in congruent and incongruent trials under different validity conditions. Results showed that incongruent trials led to longer thresholds than congruent trials in the 75%-validity condition. Furthermore, thresholds under the 100%-validity condition were not lower than those of the valid trials of other validity conditions. In other words, valid predictions did not speed up sensory processes; instead, violation of predictions slowed them down.

Nested hypothesis testing using a recursive Bayesian model suggested that the larger temporal thresholds when predictions are violated could simply be explained by the larger number of iterations needed to reach a decision without changing any internal parameters of the system (Urgen & Boyaci, 2021).

Figure 2 shows the simulation results and the model predictions for all trial types and validity conditions. The model predictions agree well with the empirical findings: The average temporal threshold is larger in the 75%-validity condition when the cues are invalid compared to when they are valid (*t*(7) = 2.91, *p* = 0.023, Cohen’s *d* = 1.03), indicating that the model also needs a longer time to detect the location of the target image when predictions are violated. Furthermore, consistent with the empirical results, there are no other significant differences between conditions. (2 (trial type: congruent, incongruent) x 2 (validity: 75%, 50%) repeated measures ANOVA results: significant main effect of congruency, *F* (1,7) = 18.52, *p* = 0.004, *η*^2^ = .73, no main effect of validity and no interaction, *F(*1,7) = 0.004, *p* = 0.951; *F*(1,7) = 0.31, *p* = 0.593; no difference between the congruent and incongruent trials in 50%-validity condition, *t*(7) = 1.4, *p* = 0.205, as well as no differences between the 100% validity condition and the congruent trials of 50%, *t*(7) = —1.67, *p* = 0.138, and 75% validity conditions, *t*(7) = —0.64, *p* = 0.54).

**Figure 2:**
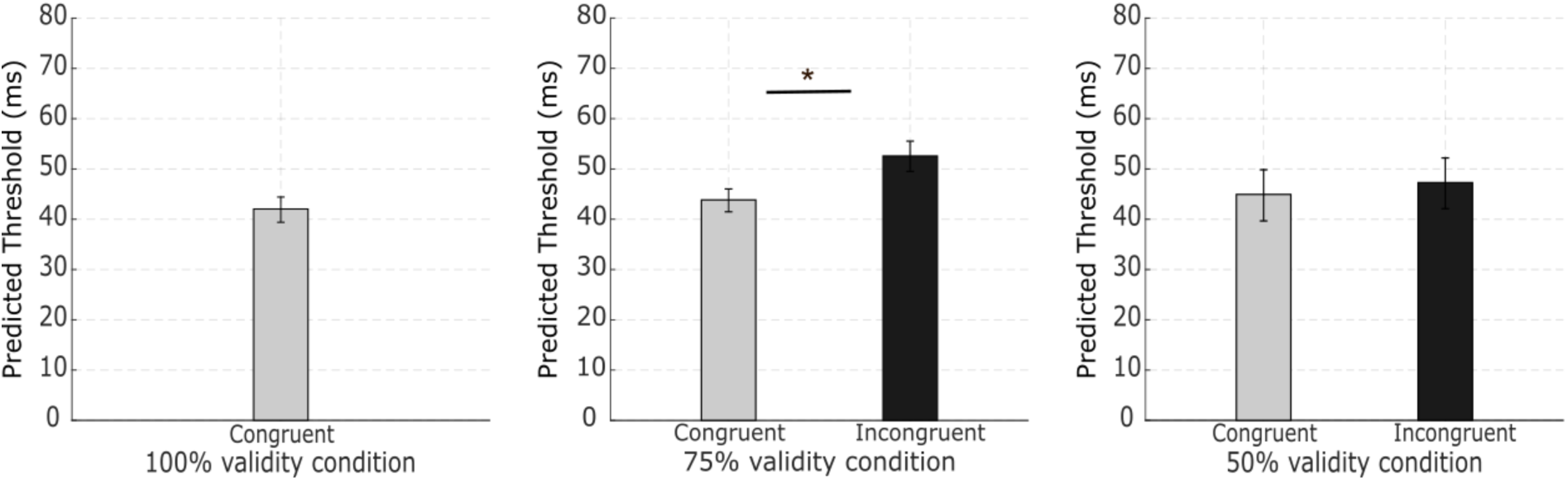
Model simulations. Group average of the predicted temporal thresholds. Consistent with the empirical results, the model predicts a higher threshold under the incongruent condition than the congruent condition when the cue validity is 75%, and no other significant differences. (Error bars: twice the standard error of the mean, *: p < 0.05)

The model simulations suggest that the differences in temporal thresholds can be explained by the larger number of iterations required when predictions are violated, as was also found with the recursive Bayesian model in Urgen and Boyaci (2021). Inspecting the unit activity time courses provides a conceptual insight for this. Figure 3 shows Layer 3 unit activities in congruent and incongruent trials (the pattern was consistent across all layers, but for clarity, only the decision layer, i.e., Layer 3, activities are shown here). The trial is from the 75%-validity condition, in which a face cue is presented. Thus, because of the prior, at the beginning of the trial, face units usually respond more strongly than other units. However, the responses dynamically change throughout the trial. Specifically, in the congruent trials (i.e., when the presented image is a face), face units remain strongly active from the onset, and the decision criterion based on the summed activity of face and house units is reached with relatively few iterations. Whereas, in the incongruent trials (i.e., when the presented image is a house), face unit responses decrease while house unit responses gradually increase, keeping their summed activity relatively low at first. To reach the decision criterion, the system needs to perform more computational iterations, thus the temporal thresholds increase.

**Figure 3:**
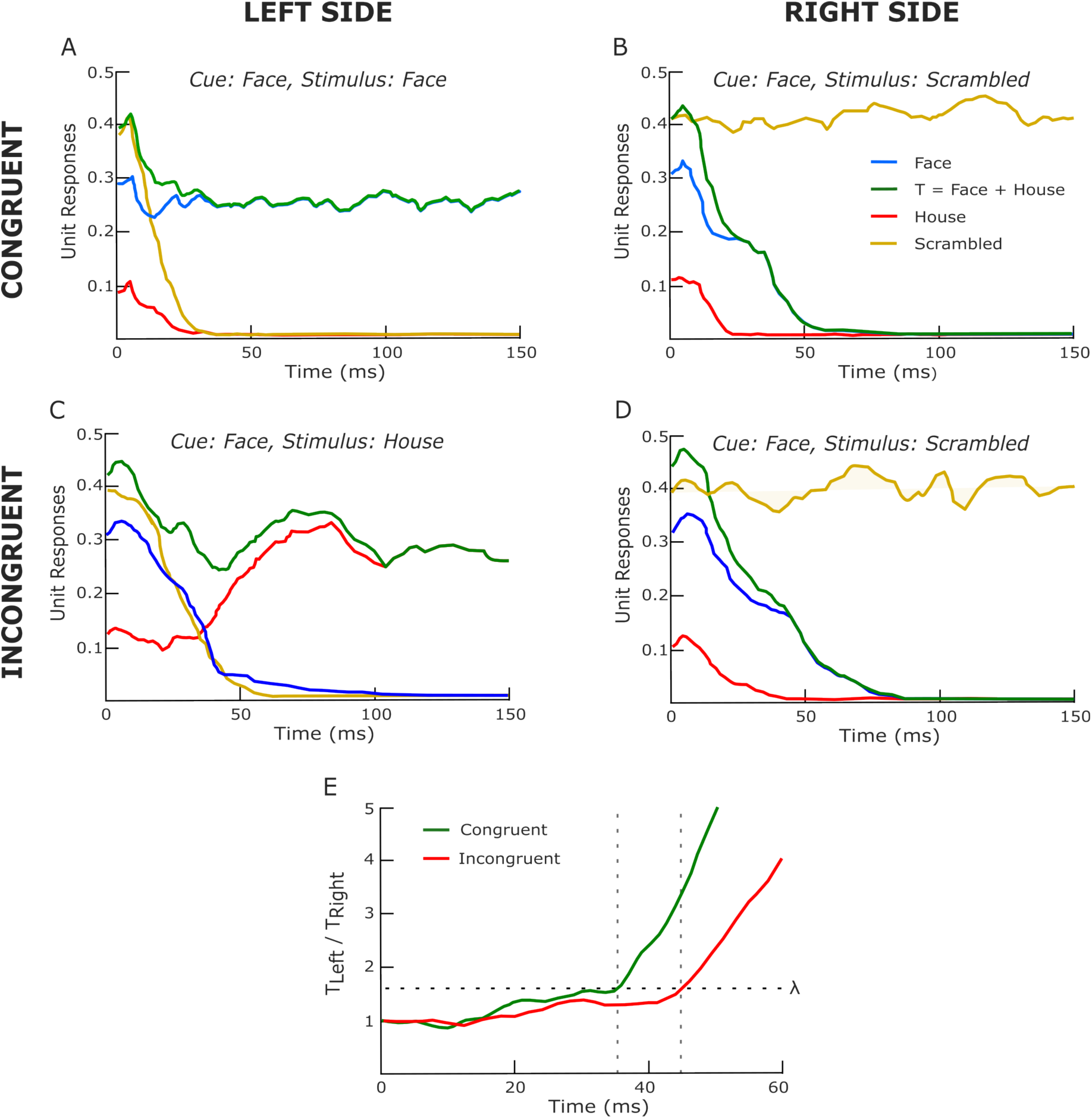
Model unit activities in sample trials. These two trials start with a *face cue* under the 75%-validity condition. The top two plots show Layer 3 unit responses to images presented on the left and on the right on a congruent trial, i.e., when the target image is a face. The lower two plots show unit responses in an incongruent trial, i.e., when the target is a house. In these simulated trials, the intact images are presented on the left. Colored lines show face, house, and scrambled unit responses, as well as the summed responses of house and face units, on which the system bases its decision. Notice the difference between the dynamic change of unit responses under the congruent and incongruent conditions. The plot at the bottom shows the time course of the ratio of the sum of face and house unit responses to the left and right images. The decision criterion, *λ*, marked by the horizontal dashed line, is reached earlier in the congruent trial compared to the incongruent trial, which is consistent with the lower perceptual thresholds when the predictions are met.

### Neural Data Simulations

To examine whether our proposed model can capture existing neuronal findings, we simulated three fMRI experiments from the literature using the optimized parameter values from our behavioral model fit above (See Supplementary Material). Stated differently, we ran simulations of the model as if our participants attended these fMRI experiments. This method was chosen for validation instead of separate model fitting to avoid possible issues such as overfitting, and it offers a stronger test for robustness and generalizability.

Studies used for the simulations were selected based on several criteria previously underlined in the field (Feuerriegel et al., 2021): *(i)* using probabilistic cueing designs with a neutral expectation/prediction condition, *(ii)* keeping the cues task-irrelevant and thereby manipulating mainly ‘prediction’ rather than ‘attention’ and/or controlling for motor response preparation, *(iii)* having cortical regions of interest (ROI) at different levels of the visual hierarchy (i.e., from V1 to FFA and MT+, up to DLPFC). All studies included the same cue validity levels: a 75%-valid and 25%-invalid predictive cue condition and a nonpredictive 50%-50% condition.

#### Egner et al. (2010)

We first simulated the experimental paradigm of Egner et al. (2010), which was used to examine FFA responses to house and face stimuli during an inverted target detection task. As in all studies covered, a probabilistic cuing paradigm was used with a 75%-valid and 25%-invalid predictive cue condition and a nonpredictive 50%-50% condition.

To simulate FFA BOLD responses, the activity of feature-1 (face) units in *Layer 2* was used, assuming that they correspond to the face-selective neurons in that area. We simulated BOLD responses under three types of trials: high face prediction trials, where a face image is predicted under the 75% validity condition; low face prediction trials, where a house image is predicted under the 75% validity condition (i.e., face prediction is 25%); and neutral trials, where the face and house images were equally likely (50% condition).

Figure 4 shows the simulation results alongside the empirical BOLD responses. We found a good agreement between the simulated and empirical results after repeating the same analyses on the simulated data as the authors did on the empirical data in the original study (Egner et al., 2010). Consistent with the empirical data analysis, 2 (stimulus type: face, house) x 3 (prediction condition: predicted, neutral, unpredicted) repeated measures ANOVA test revealed a main effect of stimulus type (*F*(1, 7) = 12.19, *p* = .01), with larger responses to face stimuli than house stimuli, no main effect of prediction (*F*(2, 14) = 1.17, *p* = 0.44), and a significant interaction between stimulus type and prediction (*F*(2, 14) = 3.81, *p* = .048). In the original study, the authors found that the difference in activity between face and house stimuli decreased with face prediction. Specifically, the difference was significant under the 25% and 50% validity conditions, but not under the 75% validity condition. Our simulation results are largely consistent with this: we found no difference between the activity for faces and houses under the 75% validity condition (*t*(7) = 0.95, *p* = .37), significant difference under the 50% validity condition (*t*(7) = 4.41, *p* = .0031), and marginally significant difference under the 25% validity condition (*t*(7) = 1.87, *p* = .1).

**Figure 4:**
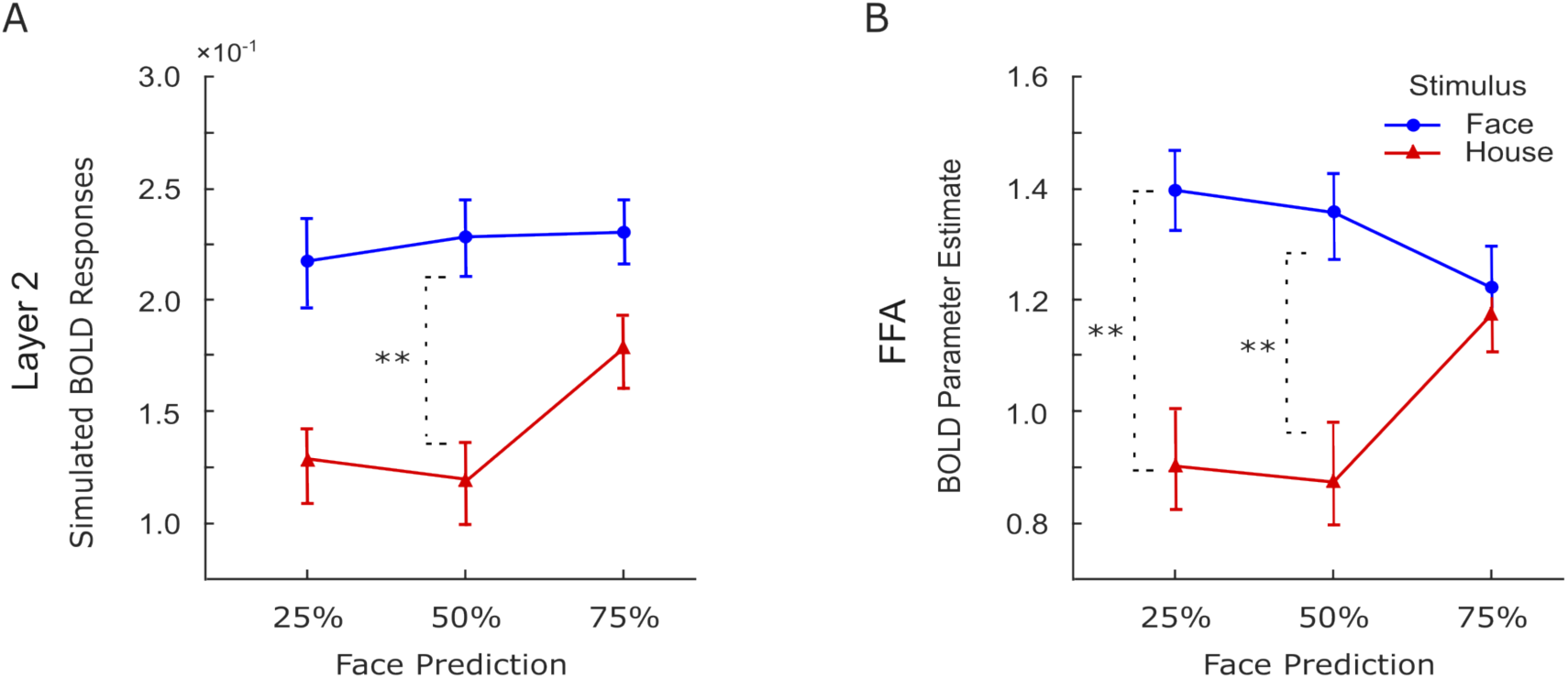
Egner et al. (2010). *A*. Group average (N = 8) of simulated BOLD responses generated with Egner et al. (2010)’s experimental procedure, using activity of Layer 2 face units. Blue circles represent responses to face stimuli, and red triangles to house stimuli. Responses to faces are overall larger than responses to houses, particularly significant in the 50% prediction condition, and marginally significant in the 25% condition. *B*. Empirical BOLD fMRI responses from FFA in the original study (adapted from Egner et al. (2010)). Faces are linked with larger FFA activity than houses, especially significant in the 25% and 50% prediction conditions. (Error bars: standard error of the mean, *p < 0.05, **p < 0.01)

#### Kok et al. (2011)

Next, we simulated BOLD responses using the protocol of Kok et al. (2011), where the authors examined the influence of cue validity on V1 BOLD responses to oriented gratings (two orientations: vertical and horizontal). Because V1 is a relatively early level area in the visual processing hierarchy, we used the summed activity of Layer 1 feature-1 and feature-2 units for the simulations. Figure 5 shows the results. Once again, the results show that the model simulations agree well with the empirical data. Kok et al. (2011) found larger BOLD responses to predicted stimuli compared to unpredicted stimuli, and the responses to the “neutral” condition were situated between the two. The simulated BOLD responses agree with this pattern: repeated measures ANOVA test revealed a significant difference across prediction conditions (*F*(2, 14) = 10.41, *p* = .002), where Bonferroni-corrected post hoc t-tests showed larger BOLD responses for the predicted condition compared to neutral (*t*(7) = 3.2, *p* = .015) and unpredicted conditions (*t*(7) = 7.3, *p* < .001).

**Figure 5:**
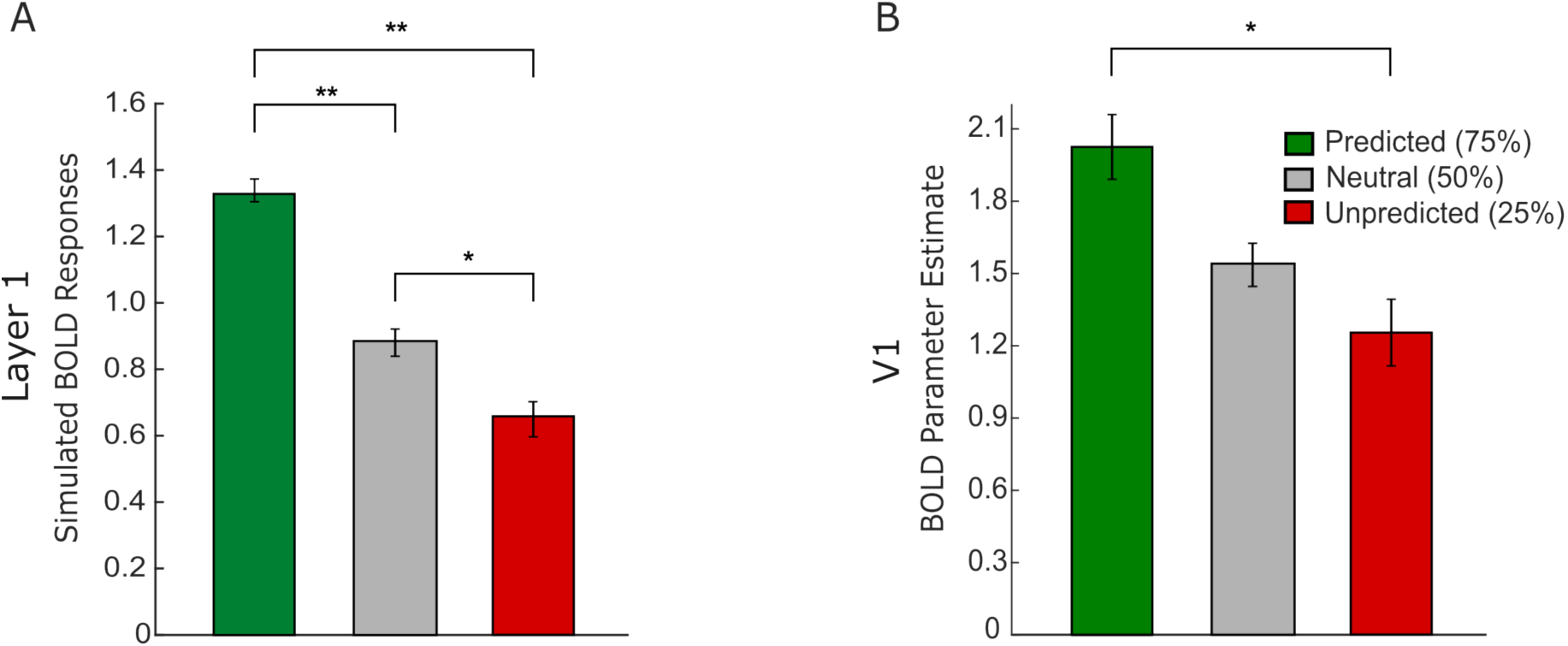
Kok et al. (2011). Vertical and horizontal gratings were used in the original design. *A*. Simulated BOLD responses using the sum of the activity of feature-1 and feature-2 units in *Layer 1*. Green and red bars represent predicted and unpredicted trials under the 75% validity condition, and the gray bar represents neutral trials (50% validity). Predicted stimuli evoked larger responses than unpredicted and neutral ones, indicating facilitation by prediction (also known as expectation facilitation). *B*. Empirical BOLD fMRI responses from V1 in the original study (adapted from Kok et al. (2011); Figure 3A, attended side). In line with the simulation results, predicted stimuli were associated with larger BOLD responses than unpredicted ones. (Error bars: standard error of the mean, \**p* < 0.05, \*\**p* < 0.01).

#### Rahnev et al. (2011)

Rahnev et al. (2011) used moving dot stimuli within a motion direction discrimination task (expanding versus contracting) and focused on hMT+, intraparietal sulcus (IPS), and dorsolateral prefrontal cortex (DLPFC) as ROIs. The effect of cued predictions diverged across regions: there was no modulation within hMT+, and higher activity with predictive cues (both valid/predicted and invalid/unpredicted) than non-predictive cues (neutral 50%-50% validity) in IPS and DLPFC (Figure 6*B-D-F*).

**Figure 6:**
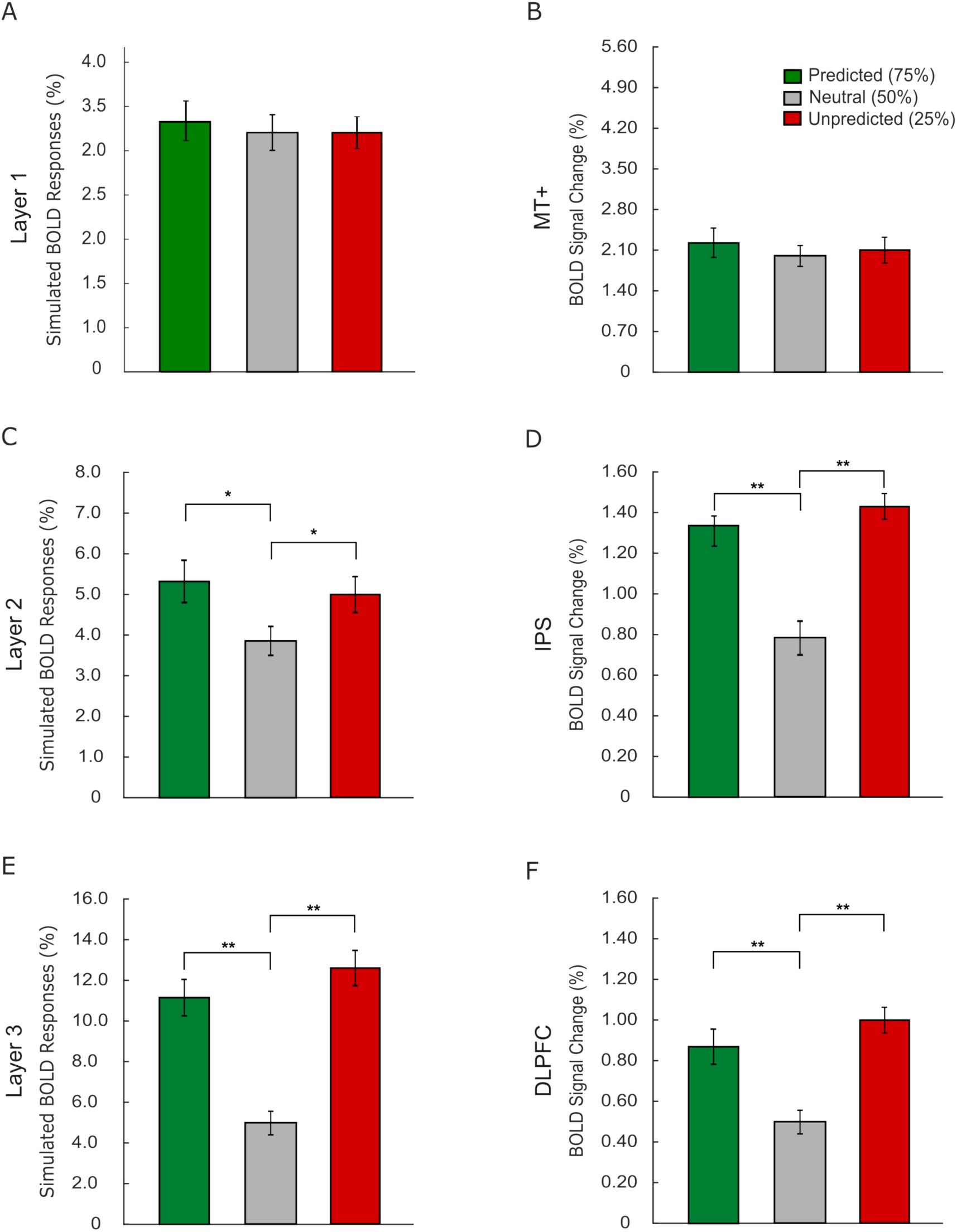
Rahnev et al. (2011). Participants discriminated between contracting and expanding moving dots in the original study. *A-C-E*. Simulated BOLD responses, obtained by the sum of the activity of feature-1 and feature-2 units from Layer 1 up to Layer 3. Green and red bars represent predicted (expected) and unpredicted (unexpected) trials under the 75% validity condition, and the gray bar represents neutral trials (50% validity). *B-D-F*. Empirical BOLD fMRI responses from hMT+, IPS, and DLPFC in the original study (adapted from Rahnev et al., 2011). (Error bars: standard error of the mean, \**p* < 0.05, \*\**p* < 0.01).

To compare simulated BOLD responses with the empirical data, we assumed a correspondence between Layer 1 of the model and hMT+, Layer 2 and IPS, and Layer 3 and DLPFC. We start by explaining our reasoning behind these assumptions. hMT+ is involved in motion processing and is placed relatively early in the visual hierarchy. It receives direct driving input from subcortical structures, as well as from V1 (Sincich et al., 2004). Thus, model Layer 1 corresponds reasonably well to hMT+. IPS, on the other hand, is implicated in higher-order motion processing such as guiding selective attention to motion and location (Gillebert et al., 2011), and maintenance of working memory related to motion (Todd & Marois, 2004; Xu & Chun, 2006). Further, it receives input from hMT+ (Greenberg et al., 2012). Thus, it corresponds well to Layer 2 of the model. Finally, DLPFC is implicated in cognitive control and conflict adaptation in general (Egner & Hirsch, 2005; Gbadeyan et al., 2016), and higher-order functions like motion sequence learning and response planning in motion processing (Badoud et al., 2016; Cieslik et al., 2013). Recently, both correlational fMRI (Kobayashi & Hsu, 2017) and causal tDCS studies (Schulreich & Schwabe, 2021) also suggested that DLPFC may be involved in belief updating under uncertainty, wherein both the probabilities of events, their reliability, and associated reward/costs are integrated and upkept. Therefore, it corresponds well to Layer 3 (the final layer) of our model, which provides the readout for the final decision.

Figure 6 shows the simulated and empirical results. In simulations, we used the sum of the activity of feature-1 and feature-2 units in each layer in predicted, unpredicted, and neutral trials. For cross-comparability, we also scaled each HRF-convolved predictor to unit height and converted condition betas to percent signal change. Results show that the empirical data and model simulations align well at all levels of the hierarchy. Neither the hMT+ activity nor model Layer 1 activity is significantly modulated by differing prediction conditions (model: *F*(2, 14) = 0.55, *p* = .59). IPS activity and Layer 2 unit responses, in contrast, both show significantly larger BOLD response for predictive cue conditions (predicted and unpredicted) compared to the neutral condition (model: *F*(2, 14) = 4.47, *p* = 0.032; predicted vs. neutral: *t*(7) = 2.84, *p* = 0.025, unpredicted vs. neutral: *t*(7) = 2.57, *p* = 0.037). DLPFC and model Layer 3 responses are similar to those found in the previous level, with a sharper difference between predictive and nonpredictive conditions (model: *F*(2, 14) = 7.78, *p* = 0.005; predicted vs. neutral: *t*(7)= 3.69, *p* = 0.008, unpredicted vs. neutral: *t*(7) = 4.12, *p* = 0.004). Thus, overall, model simulations and empirical findings match at all examined brain regions.

## Discussion

Our results show that the proposed model can successfully explain the behavioral and neuronal effects of predictions and approximate Bayesian inference. Two important aspects of the model simulations are worth reiterating. Firstly, we optimized the model parameters using the behavioral data and simulated the fMRI data using those parameters. We did not fit the model to the fMRI data. In a sense, we simulated fMRI data using the protocols of the selected fMRI studies as if they were collected from the participants of the behavioral experiment.

Secondly, the selected fMRI studies were quite diverse, focusing on widely varied regions within the visual hierarchy, from V1 to FFA and hMT+ up to DLPFC, covering both the dorsal and the ventral pathways. Furthermore, they used different sets of stimuli (oriented gratings, faces, houses, and moving dots), whereas the behaviora experiment tested only faces and houses. It is remarkable that the model was able to capture the pattern of neuronal responses with such diversity by simply using parameters optimized with the behavioral data of a single study. Overall, this stringent and unbiased approach demonstrates that the model has great promise in terms of scope and generalizability, with relatively few assumptions made.

### Comparison with the Predictive Coding Model

One of the mathematically most developed models of cortical functioning is the predictive coding model (PC), an idea that dates back to the 1990s (D. Mumford, 1992). The model was first rigorously introduced by Rao and Ballard (1999), then put in a variational inference form by Friston (e.g., Friston, 2005). Similar to the model proposed here, the predictive coding model also posits that predictions, through top-down information flow, shape neural processes. Predictive coding models hypothesize that higher-level brain areas generate a model of the world that is relayed to lower levels, just as the model presented here. In the lower levels, the difference between sensory input and those predictions is computed, and any residual error is signaled back to higher levels to correct and refine the models of the world. According to the PC model, the effect of feedback is largely to suppress the activity in lower areas for predicted stimuli, because they receive inhibitory input from higher levels (subtraction). This is, however, not consistent with the bulk of findings in visual neuroscience: the activity may reduce but does not vanish even if a stimulus is predicted (e.g., Burns et al., 2010). Whereas our model posits the opposite, if a stimulus is predicted, units that participate in its processing do not necessarily reduce their activity; in fact, they may become activated even in the absence or prior to the presentation of a physical stimulus that is predicted.

Specifically, in our model simulations (i.e., cortical model unit responses, Figure 3), we observe an early prior-driven bias in favor of the predicted-object category (i.e., faces): at trial onset, those predicted-object-selective units dominate. In valid trials, this bias aligns with the sensory evidence, sustaining strong predicted-object activity and enabling the decision bound to be reached in fewer iterations. In invalid (unpredicted, i.e., house) trials, the system reweights over time: predicted-object activity decays while unpredicted-object activity gradually increases, keeping their sum initially low and thus requiring more computational iterations to reach the bound. This pattern is consistent with dynamic inference frameworks (e.g., Heeger, 2017) and with selective, excitatory feedback plus feedforward adaptation (e.g., Westerberg & Roelfsema, 2025), and it is also supported by many findings in visual neuroscience literature (e.g., Kok et al., 2014, 2017).

Although PC has been very successful in compressing information transmission in engineering applications, its biological plausibility remains uncertain. The implementation of PC models in the brain requires *representation units* and at least one type of *error units*. Often, one type of error unit is not sufficient, and situations where an expected stimulus is not received are addressed by introducing “absence error units”. Because errors may need to be computed between different features (e.g., houses and faces), the feature selectivity of the error units must also be accounted for, which further increases the number of required error unit types owing to the large number of possible combinations to compare. There are several studies in the literature that suggest the possible microcircuitry required to satisfy the demands of the PC model (Bastos et al., 2012; Shipp, 2016). These efforts have yet to outline a biologically plausible mechanism (Millidge et al., 2022).

In contrast to circuit-level predictive coding accounts that posit distinct error and representation units, our framework is formulated at the computational level that emphasizes functional interactions among neural populations. Instead of requiring additional population types that are not clearly supported by current physiological evidence, our model builds upon established cortical circuitry to support predictive processing. Specifically, the model tested here requires feature-tuned neurons receiving feedback and feedforward input, as well as making within-area lateral interactions, all of which are known to exist based on our current knowledge of the early visual system. For example, cortical layer 4 of V1 receives both feedforward input from LGN and feedback from higher areas either directly or through superficial and deep layers (Vanni et al., 2020).

Our fMRI simulations show that the model predictions are consistent with empirical findings. For example, according to the model, increased fMRI activity to unpredicted stimuli could be caused by prolonged computations (J. A. Mumford et al., 2023), as well as the concurrent activity of the population of neurons tuned to the predicted and unpredicted features. Results of several neuroimaging studies, nevertheless, appear to be consistent with, and thus taken as supporting evidence for, PC models (although see Alink & Blank, 2021; Feuerriegel, 2024; Feuerriegel et al., 2021). Specifically, increased neural response to unpredicted stimuli (compared to predicted stimuli) is interpreted to reflect the signals of feature-selective error units (Kok et al., 2011; Summerfield & Egner, 2009). Some work, however, interprets findings as reflecting decreased neural response to predicted stimuli, introducing the *expectation suppression* effect (Richter & de Lange, 2019; Summerfield & Koechlin, 2008).

Interestingly, depending on the findings, authors of the studies appear to claim that PC models explain either type of results. This is because PC models are very flexible, which is not surprising given that they are based on variational inference algorithms with diverse possible neural implementations. Perhaps because of this, PC models have typically not been subjected to pre-planned, direct, rigorous empirical validation using behavioral and neural data, and instead have often functioned as post hoc tools for interpreting results qualitatively. The variability in the neural patterns may also suggest that several results that are attributed to prediction may actually arise under different contextual and computational conditions, and involve additional mechanisms such as novelty response, attentional modulation, or adaptation rather than only prediction-related influences (see Feuerriegel et al., 2021). Consistent with this, the expectation suppression effect has been predominantly reported in a subset of paradigms that involve statistical learning designs with extensive training periods, but short-term probabilistic/predictive cueing studies provided inconsistent evidence for such an effect (den Ouden et al., 2025; Feuerriegel, 2024; Feuerriegel et al., 2021). We argue that our simulations capture the context in which predictive feedback strengthens the representations of expected inputs in the visual system, and our proposed model offers a more principled, parsimonious, and testable explanation for the neuroimaging findings compared to PC models.

An implementation of the PC model, with an appropriate read-out rule, could, in principle, yield predictions that align with the behavioral and fMRI data tested here. However, we do not undertake such simulations because our primary goal is not to quantitatively compare different models, but rather to demonstrate that an alternative model can make successful predictions. Furthermore, as explained before, PC is extremely flexible to the point that it can fit almost any data provided to it (Millidge et al., 2022), thus rendering any model comparison difficult to interpret. Our study highlights the importance of employing and testing predictive processing models against empirical data, with the hope of inspiring further exploration and evaluation of the unique constructs proposed by various predictive processing models. Future work could extend the current approach by testing the model in experiments designed for direct model comparison by systematically manipulating stimulus features, sensory uncertainty, and prediction strength. Incorporating temporal dynamics and laminar-specific data could further constrain the neural implementation.

## Conclusion

In sum, we present a parsimonious predictive processing model of the functioning brain, which can explain the effects of predictions, expectations, and prior knowledge on behavior and neural activity. It offers a possible mechanism underpinning Bayesian inference in the brain and forms a rigorous and testable link between behavioral and neural responses. The model is biologically plausible in that all the building blocks required by the model are already known to exist in the neural system. By bridging behavioral and neural data within a single framework, our approach advances understanding of how predictions influence sensory processing and offers a principled, parsimonious predictive processing model.

## Data and Code Availability

The data and code used in the study will be made available to readers.

## Acknowledgements

This work was funded by a grant from the Turkish National Scientific and Technological Council (TÜ BİTAK 217K163) awarded to HB. We thank Katja Doerschner and Burcu Aysen Urgen for their valuable comments on an earlier version of the manuscript.

## Author Contributions

BMU, HIB, and HB designed the research, performed the research, analyzed the data, and wrote the paper.

## Competing Interests

The authors declare no competing interests.

## Notes

### Competing Interest Statement

The authors have declared no competing interest.

### Summary of Updates

This revised version extends the original manuscript beyond behavioral modeling by incorporating simulations of neural responses. Using parameters optimized from behavioral data we evaluate the model's predictions against published fMRI findings showing that the same framework accounts for both behavioral and neural effects of prediction.

## References

Alink, A., & Blank, H. (2021). Can expectation suppression be explained by reduced attention to predictable stimuli? NeuroImage, 231, 117824.

Badoud, S., Borgognon, S., Cottet, J., Schmidlin, E., Kaeser, M., & Rouiller, E. M. (2016). Effects of dorsolateral prefrontal cortex lesion on motor habit and performance assessed with manual grasping and control of force in macaque monkeys. Brain Structure and Function, 222 (7), 3299–3321. 10.1007/s00429-016-1268-z

Bastos, A. M., Usrey, W. M., Adams, R. A., Mangun, G. R., Fries, P., & Friston, K. J. (2012). Canonical microcircuits for predictive coding. Neuron, 76 (4), 695–711. 10.1016/j.neuron.2012.10.038

Burns, S. P., Xing, D., & Shapley, R. M. (2010). Comparisons of the Dynamics of Local Field Potential and Multiunit Activity Signals in Macaque Visual Cortex. Journal of Neuroscience, 30 (41), 13739–13749. 10.1523/JNEUROSCI.0743-10.2010

Cieslik, E. C., Zilles, K., Caspers, S., Roski, C., Kellermann, T. S., Jakobs, O., Langner, R., Laird, A. R., Fox, P. T., & Eickhoff, S. B. (2013). Is there “one” dlpfc in cognitive action control? evidence for heterogeneity from co-activation-based parcellation. Cerebral Cortex, 23 (11), 2677–2689. 10.1093/cercor/bhs256

de Lange, F. P., Heilbron, M., & Kok, P. (2018). How do expectations shape perception? Trends in cognitive sciences.

den Ouden, C., Kashyap, M., Kikkawa, M., & Feuerriegel, D. (2025). Limited evidence for probabilistic cueing effects on grating-evoked event-related potentials and orientation decoding performance. Psychophysiology, 62 (5), e70076.

Egner, T., & Hirsch, J. (2005). Cognitive control mechanisms resolve conflict through cortical amplification of task-relevant information. Nature Neuroscience, 8 (12), 1784–1790. 10.1038/nn1594

Egner, T., Monti, J. M., & Summerfield, C. (2010). Expectation and surprise determine neural population responses in the ventral visual stream. Journal of Neuroscience, 30 (49), 16601–16608.

Feuerriegel, D. (2024). Adaptation in the visual system: Networked fatigue or suppressed prediction error signalling? Cortex.

Feuerriegel, D., Vogels, R., & Kovács, G. (2021). Evaluating the evidence for expectation suppression in the visual system. Neuroscience & Biobehavioral Reviews, 126, 368–381.

Friston, K. (2005). A theory of cortical responses. Philosophical Transactions of the Royal Society of London B: Biological Sciences, 360 (1456), 815–836.

Gbadeyan, O., McMahon, K., Steinhauser, M., & Meinzer, M. (2016). Stimulation of dorsolateral prefrontal cortex enhances adaptive cognitive control: A high-definition transcranial direct current stimulation study. Journal of Neuroscience, 36 (50), 12530–12536. 10.1523/JNEUROSCI.2450-16.2016

Gillebert, C. R., Mantini, D., Thijs, V., Sunaert, S., Dupont, P., & Vandenberghe, R. (2011). Lesion evidence for the critical role of the intraparietal sulcus in spatial attention. Brain, 134 (6), 1694–1709. 10.1093/brain/awr085

Greenberg, A. S., Verstynen, T., Chiu, P. H., & Yantis, S. (2012). Visuotopic cortical connectivity underlying attention. Journal of Neuroscience, 32 (8), 2773–2782. 10.1523/JNEUROSCI.4515-11.2012

Harrison, W. J., Bays, P. M., & Rideaux, R. (2023). Neural tuning instantiates prior expectations in the human visual system. Nature Communications, 14 (1), 5320.

Heeger, D. J. (2017). Theory of cortical function. Proceedings of the National Academy of Sciences, 114 (8), 1773–1782.

Heekeren, H. R., Marrett, S., Bandettini, P. A., & Ungerleider, L. G. (2004). A general mechanism for perceptual decision-making in the human brain. Nature, 431 (7010), 859.

Kobayashi, K., & Hsu, M. (2017). Neural mechanisms of updating under reducible and irreducible uncertainty. Journal of Neuroscience, 37 (29), 6972–6982. 10.1523/JNEUROSCI.0279-17.2017

Kok, P., Failing, M. F., & de Lange, F. P. (2014). Prior Expectations Evoke Stimulus Templates in the Primary Visual Cortex. Journal of Cognitive Neuroscience, 26 (7), 1546–1554. 10.1162/jocn_a_00562

Kok, P., Mostert, P., & Lange, F. P. D. (2017). Prior expectations induce prestimulus sensory templates. Proceedings of the National Academy of Sciences of the United States of America, 114, 10473–10478. 10.1073/PNAS.1705652114/SUPPL_FILE/PNAS.201705652SI.PDF

Kok, P., Rahnev, D., Jehee, J. F., Lau, H. C., & De Lange, F. P. (2011). Attention reverses the effect of prediction in silencing sensory signals. Cerebral cortex, 22 (9), 2197–2206.

Ma, W. J., Beck, J. M., Latham, P. E., & Pouget, A. (2006). Bayesian inference with probabilistic population codes. Nature Neuroscience, 9 (11), 1432–1438. 10.1038/nn1790

Millidge, B., Seth, A. K., & Buckley, C. L. (2022). PREDICTIVE CODING: A THEORETICAL AND EXPERIMENTAL REVIEW.

Mumford, D. (1992). On the computational architecture of the neocortex. Biological cybernetics, 66 (3), 241–251.

Mumford, J. A., Bissett, P. G., Jones, H. M., Shim, S., Ali, J., Rios, H., & Poldrack, R. A. (2023). The response time paradox in functional magnetic resonance imaging analyses. Nature Human Behaviour 2023, 1–12. 10.1038/s41562-023-01760-0

Rahnev, D. A., Lau, H. C., & de Lange, F. P. (2011). Prior expectation modulates the interaction between sensory and prefrontal regions in the human brain. Journal of Neuroscience, 31 (29), 10741–10748. 10.1523/JNEUROSCI.1478-11.2011

Rao, R. P., & Ballard, D. H. (1999). Predictive coding in the visual cortex: A functional interpretation of some extra-classical receptive-field effects. Nature neuroscience, 2 (1), 79.

Richter, D., & de Lange, F. P. (2019). Statistical learning attenuates visual activity only for attended stimuli. elife, 8, e47869.

Schulreich, S., & Schwabe, L. (2021). Causal role of the dorsolateral prefrontal cortex in belief updating under uncertainty. Cerebral Cortex, 31 (1), 184–200. 10.1093/cercor/bhaa219

Shipp, S. (2016). Neural elements for predictive coding. Frontiers in psychology, 7, 1792.

Sincich, L. C., Park, K. F., Wohlgemuth, M. J., & Horton, J. C. (2004). Bypassing v1: A direct geniculate input to area mt. Nature Neuroscience, 7 (10), 1123–1128. 10.1038/nn1318

Summerfield, C., & De Lange, F. P. (2014). Expectation in perceptual decision making: Neural and computational mechanisms. Nature Reviews Neuroscience, 15 (11), 745–756.

Summerfield, C., & Egner, T. (2009). Expectation (and attention) in visual cognition. Trends in cognitive sciences, 13 (9), 403–409.

Summerfield, C., & Koechlin, E. (2008). A neural representation of prior information during perceptual inference. Neuron, 59 (2), 336–347.

Todd, J. J., & Marois, R. (2004). Capacity limit of visual short-term memory in human posterior parietal cortex. Nature, 428 (6984), 751–754. 10.1038/nature02466

Urgen, B. M., & Boyaci, H. (2021). Unmet expectations delay sensory processes. Vision Research, 181, 1–9. 10.1016/j.visres.2020.12.004

Vanni, S., Hokkanen, H., Werner, F., & Angelucci, A. (2020). Anatomy and Physiology of Macaque Visual Cortical Areas V1, V2, and V5/MT: Bases for Biologically Realistic Models. Cerebral Cortex, 30 (6), 3483–3517. 10.1093/CERCOR/BHZ322

Westerberg, J. A., & Roelfsema, P. R. (2025). Hierarchical interactions between sensory cortices defy predictive coding. Trends in Cognitive Sciences.

Xu, Y., & Chun, M. M. (2006). Dissociable neural mechanisms supporting visual short-term memory for objects. Nature, 440 (7080), 91–95. 10.1038/nature04262

