## Supplementary Material for "A recurrent neuronal model for the effect of predictions on sensory processes"

### Supplementary Tables and Figures

Supplementary Table 1: **Optimized parameters of the model for all participants.**

| subject# | $\Delta t$ | $\hat{\sigma}$ | $\lambda$ |
|---|---|---|---|
| 1 | 3.11 | 0.41 | 1.52 |
| 2 | 2.04 | 0.36 | 1.82 |
| 3 | 1.61 | 0.30 | 2.07 |
| 4 | 1.72 | 0.27 | 1.97 |
| 5 | 1.38 | 0.40 | 1.22 |
| 6 | 2.20 | 0.39 | 1.45 |
| 7 | 1.51 | 0.39 | 2.15 |
| 8 | 1.80 | 0.43 | 2.30 |

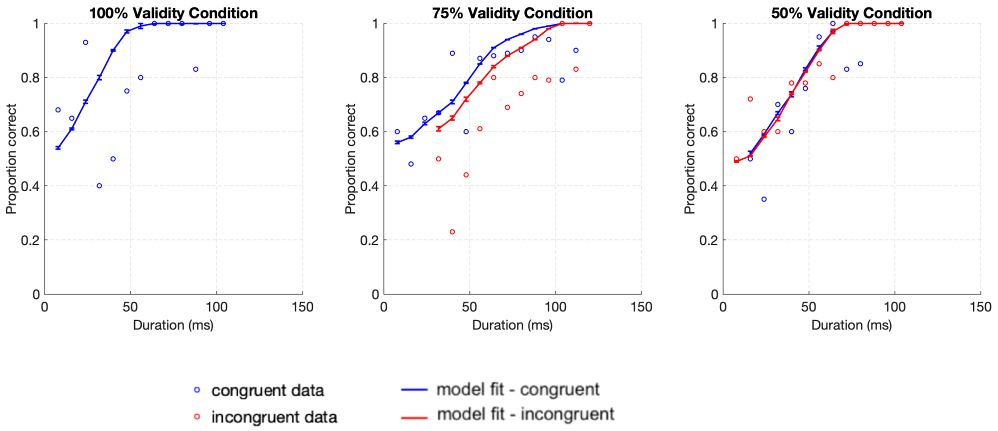


Supplementary Figure 1: **Model fits for a single participant (1).** Error bars are twice the standard error.

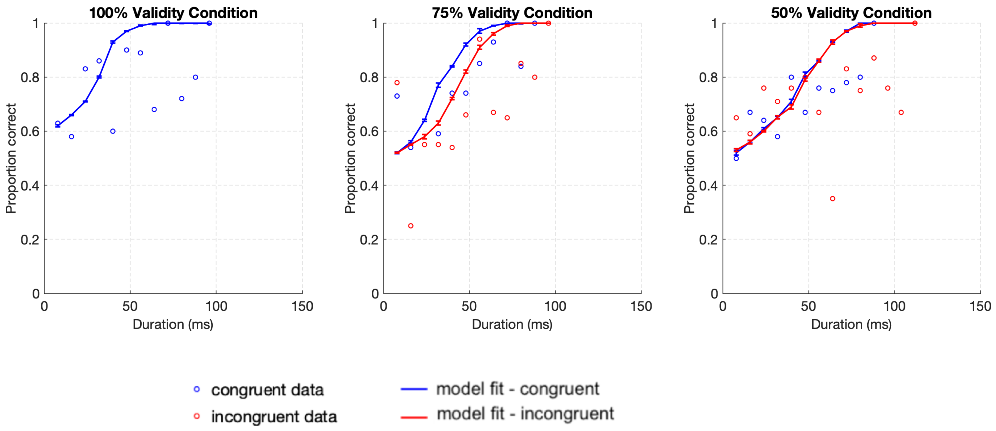


Supplementary Figure 2: **Model fits for a single participant (2).** Error bars are twice the standard error.

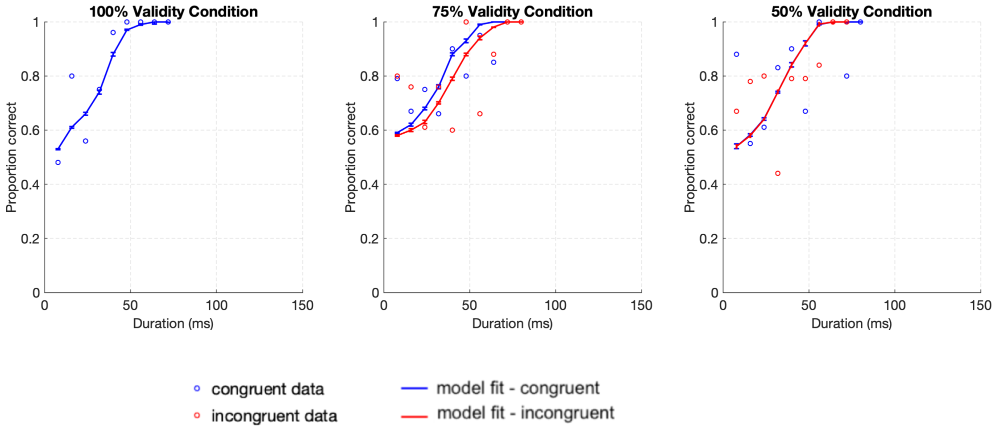


Supplementary Figure 3: **Model fits for a single participant (3).** Error bars are twice the standard error.

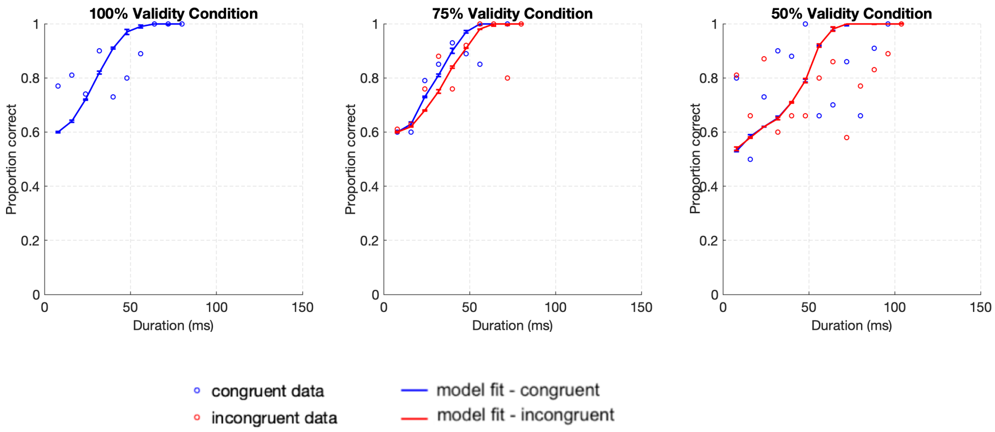


Supplementary Figure 4: **Model fits for a single participant (4).** Error bars are twice the standard error.

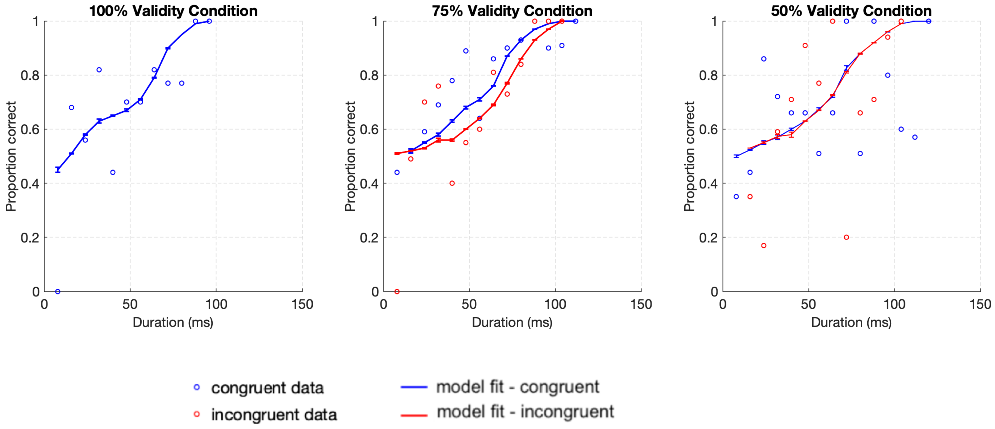


Supplementary Figure 5: **Model fits for a single participant (5).** Error bars are twice the standard error.

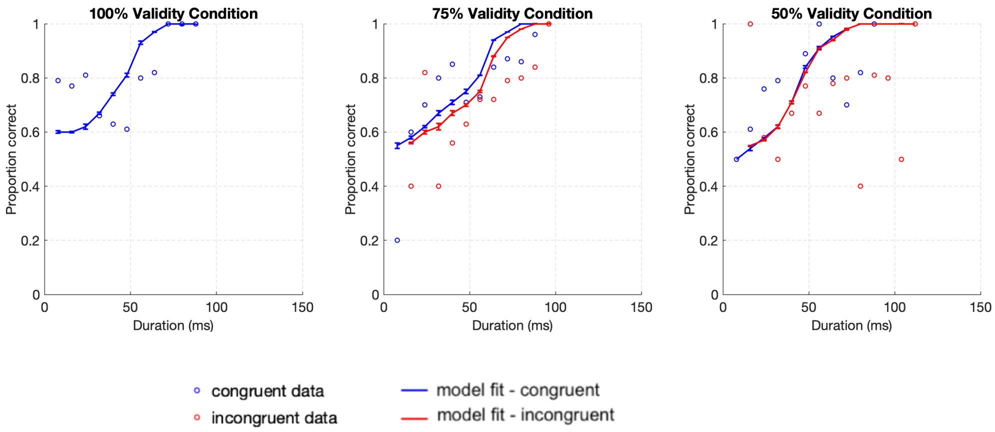


Supplementary Figure 6: **Model fits for a single participant (6).** Error bars are twice the standard error.

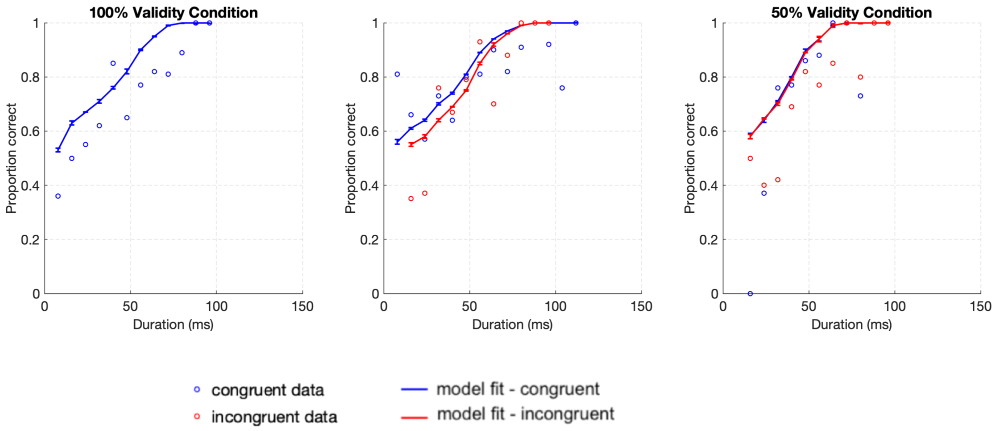


Supplementary Figure 7: **Model fits for a single participant (7).** Error bars are twice the standard error.

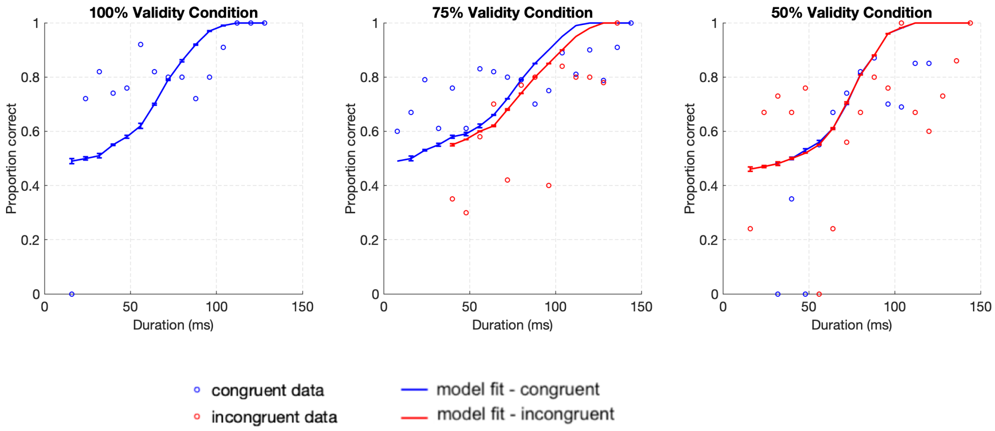


Supplementary Figure 8: **Model fits for a single participant (8).** Error bars are twice the standard error.
